# CCL19-CCR7 mediated recruitment of T cells is associated with the leishmanin skin test in individuals with prior exposure to *Leishmania* parasites

**DOI:** 10.64898/2026.02.19.706883

**Authors:** Younis Brima Musa, Saeed Mohammed Abdelrahim, Khalil Eltahir Awad Gasim, Hira L Nakhasi, Musa Ahmed Mudawi, Sreenivas Gannavaram, Abhay R Satoskar

## Abstract

A delayed-type hypersensitivity (DTH) response induced by the intradermal inoculation of leishmanin antigens is used to detect prior exposure to the protozoan parasite Leishmania. *Leishmanin* antigen preparations are an important tool in disease surveillance studies in endemic areas. Commercial scale leishmanin antigens are being developed for wider deployment in assessing vaccine efficacy and latent infections in the field. Previous studies with leishmanin induced DTH response were limited to analysis of PBMCs. To investigate the mediators of DTH response, we performed spatial transcriptomic analysis of the DTH skin biopsies obtained from a *Leishmania* endemic foci. Compared to healthy skin biopsy, macrophages and T cells, and IL-16 and TNF that attract CD4^+^ T cells, were elevated in Langerhans cells in the DTH biopsies. Both IFN-γ and its receptors were similarly elevated in the DTH biopsies. Chemokines CCL2, CCL5, CCL8 and CCL19, and the corresponding receptors were elevated in DTH biopsies, with CCL19-CCR7 as the most salient interaction in our Cellchat analysis. These data reveal biomarkers of DTH response following leishmanin inoculation and enable appropriate formulation and reintroduction of leishmanin skin test antigens for disease surveillance.

## Introduction

An estimated 700,000 to 1 million new cases annually of all forms of human leishmaniasis including visceral, cutaneous, and mucocutaneous forms occur worldwide (https://www.who.int/news-room/fact-sheets/detail/leishmaniasis). Current surveillance systems do not fully capture the burden as the disease spreads into new geographic areas including previously non-endemic zones due to climate change, urbanization, and migration (Cosma 2024). Serological assays such as the rK39 rapid diagnostic test have several limitations for VL surveillance studies in East Africa where it showed lower sensitivity (60-85%) in Ethiopia, Sudan, and Kenya, compared to the Indian subcontinent (>90%, Zijlstra 2001; Boelaert 2008). Serological tests can produce false positives in patients with other common endemic diseases including malaria, tuberculosis, trypanosomiasis, and other parasitic infections. More importantly, rK39 has limited ability to detect asymptomatic or subclinical infections, which are crucial for understanding true disease burden and transmission dynamics in surveillance studies.

In contrast to serological tests, the delayed-type hypersensitivity (DTH) reaction mediated primarily by the activity of Th1-type CD4^+^ T cells that recognize *Leishmania* antigens and produce IFN-γ is better suited to detect cured or subclinical infections. Studies emphasized positive LST correlates with protection and past exposure, not active parasitemia. A DTH positive reaction is associated with long-lasting protective immunity against reinfection, most notably for VL (Carstens-Kass2021). DTH-negative but seropositive individuals are more likely to progress to active visceral leishmaniasis (Sacks 2002; Zijlstra 2016, Pacheco-Fernandez 2025). Thus, DTH reactions following inoculation with leishmanin antigens have been employed for surveillance of CL (Osman 2021) and VL in Sudan (Zijlstra 1994; Sharief 2019). Similarly, transmission risk of American CL was assessed using DTH reactions in Brazil (Aston 1970; Souza 1992). There is currently no WHO-prequalified, GMP-grade leishmanin which has curtailed field use and cross-study comparability is not possible. Longitudinal studies of VL in Bangladesh assessing the potency of leishmanin antigens reported a decline in sensitivity emphasizing the need for better production, standardization of sensitivity, potency, and stability of leishmanin antigens (Bern 2006).

Our current understanding of the mediators of DTH reactions is primarily derived from studies of Tuberculin skin test. During the effector phase where upon re-exposure to antigen (e.g., intradermal injection of PPD in tuberculin test), memory Th1 cells recognize antigen-MHC complex on local APCs, release cytokines such as IFN-γ, and TNF-α which activate macrophages, endothelial cells, recruit monocytes/neutrophils/lymphocytes resulting in a visible skin induration 24-72 hours following inoculation (Huebner, 1993; Vukmanovic-Stejic, 2006). Following non-specific neutrophil infiltration during the initial phase (Platt 1983), CD4^+^ and CD8+ T cells, activated macrophage infiltrate the site of inoculation (Poulter 1982). Recruitment of T cells is supported by early pro-inflammatory cytokine release (IFN-γ, TNF-α, lymphotoxin) that stimulates endothelial adhesion molecule expression (E-selectin, ICAM-1, VCAM-1) and increases vascular permeability (Gibbs 1984). In addition, FOXP3+ Tregs were also observed to locally proliferate within human skin during DTH reactions, and elevated frequencies of Tregs were found in the skin lesions of contact dermatitis and resolving fixed drug eruptions (Teraki and Shiohara, 2003; Vukmanovic-Stejic et al., 2008).

Studies analyzing skin resident immune cell populations in inflammatory reactions are limited. DTH reactions from Mantoux and allergen patch tests remain the commonly used methods to investigate the skin immunity (Vukmanovic-Stejic et al., 2008) and allergen patch tests (Spergel 2005). These tests produce qualitative readouts of the response but do not provide mechanistic insights into the phenotypic and functional characteristics of the cell infiltrate. Such understanding of the coordinated activities of cytokine/chemokine signaling networks among the cells in the DTH reaction site would help produce highly optimized leishmanin antigen preparations that currently suffer from lack of standardization, intra-study comparability, and regulatory compliance.

To address this gap, we have undertaken spatial transcriptomic analysis of the archived FFPE skin biopsies of positive DTH reaction site from subjects with no known history of VL in Sudan as part of surveillance. Our data allowed us to reconstruct the cellular composition, chemokine/cytokine signaling at the DTH site. Our bioinformatic analysis illuminates the mediators of DTH reaction in not only leishmanin skin tests and may also have wider applicability to skin inflammatory reactions.

## Methods

### Description of the biopsy material

Archived formalin-fixed paraffin embedded skin biopsies originally collected under an IRB with informed consent from the study participants were used in the current analysis. All the skin biopsies are from participants with no history of visceral leishmaniasis. The skin biopsies were collected from the indurated sites of LST inoculation. The LST antigens used in the study were obtained from the Pasteur Institute, Iran. To identify individuals with asymptomatic leishmania infection, residents in VL -endemic area who are reactive to LST and with no past history of VL were included.

### Tissue Preparation for Visium CytAssist instrument

Formalin-fixed paraffin embedded (FFPE) tissue blocks for the six biopsy tissues (DTH, n=5 and healthy control biopsy from living donor n=1) were sectioned a x-micron thickness. One best section was placed in the center of a standard glass slide (frosted) and stained with hematoxylin and eosin, using standard protocol, and coverslipped. The slides were devoid of any patient identifiers and were labelled just with serial numbers that were linked to the individual patient. Spatial transcriptomic profile data capture was performed at the Ohio State University Genomics Core Laboratory, Columbus. After uncoverslipping the slides, the tissue is then de-crosslinked to release mRNA that was sequestered by formalin fixation. Human whole transcriptome probe panels, consisting of a pair of specific probes for each targeted gene, are added to the tissue. These probe pairs hybridize to their gene target and are then ligated to one another. The spatially barcoded, ligated probe products are released from the slide and PCR amplified. The probe products are further processed to generate an NGS-ready library. The Visium Spatial Gene Expression library is sequenced using standard short-read sequencers. Using license free software from 10X Genomics, data is processed and visualized using Space Ranger analysis pipelines and Loupe Browser visualization software.

### Data capture using 10x Genomics software

During the Visium workflow, two main data types are captured: a tissue image and sequencing data. For Visium FFPE with facilitated probe transfer using the Visium CytAssist instrument and third image is captured by the CytAssist to provide spatial orientation of the data. 10x Genomics provides two software tools to process and visualize these Visium data types, Space Ranger and Loupe Browser. Space Ranger processes the input file types to align the Visium sequencing data with the image. Each Spatial Barcode with the associated UMIs captured during the Visium workflow is assigned a spatial location in the tissue image. Space Ranger produces a variety of output files that can be used in Loupe Browser or third-party tools to visualize and apply spatial analysis methods to the data.

### Curation of custom reference for alignment

For the alignment step, genomic sequences and .gtf files for *Homo Sapiens* (assembly GRCh38) were merged with *Leishmania donovani* (version GCA_000009645.1) to create a custom reference. The annotation information for *L. donovani* parasite was appended to the end of the human annotation file. The merged genome and annotation custom reference was then processed by spaceranger/cellranger mkgtf and then spaceranger/cellranger mkref (version 2.1.0), using the default parameters, to create the appropriate genome index files. We aligned all libraries to the custom curated reference.

### Read processing

Base calling was performed using spaceranger mkfastq with default parameters. The sequenced reads for each sample were processed separately using spaceranger count against the host/parasite merged genome and the corresponding merged annotation gtf file.

### Spatial transcriptomics data analysis with Seurat

Count matrices produced by spaceranger count were processed using the R package Seurat (v4.4, Hao et al., 2023). We filtered out any cells with low UMI count, very few expressed genes or excessive mitochondrial gene load (indicative of dead/dying cells). scDblFinder was used to remove UMI doublets from the aggregate dataset (Germain et al, 2021). Individual samples were then integrated using Seurat’s SCTransform specific integration workflow. The cells were clustered by applying the K-nearest neighbors (KNN) graph based on PCA reduced space, followed by Louvain’s algorithm available in Seurat (v4.4). The cells were projected to a two-dimensional space using the Uniform Manifold Approximation and Projection (UMAP) dimensionality reduction technique. Annotation of computationally predicted clusters to biological cell type was performed manually using a combination of the following databases: CellMarker, CellMarker 2.0, panglaoDB, czscience. Gene cluster markers of each population were detected by identifying significant differentially expressed genes between one population and the rest of the cells using Wilcoxon test, available in Seurat package (v4.4), including only genes expressed in at least 25% of the cells of either group. For differential gene expression analysis we utilized both DESEQ2 and MAST algorithms.

### Cell Communication Network (CCN) analysis using CellChat

To investigate cell communication patterns across the identified cell types we used CellChat (Jin et al., 2021). In order to achieve compatibility we treated the spatial RNAseq matrices as single cell RNAseq matrices and loaded the data using Read10X function instead of Load10X_Spatial. Computational clusters were confirmed to match the exact clusters produced when using the Load10X_Spatial results function. We focused on secreted signaling communication database composed from thousands of ligand-receptor combinations tailored for single cell RNA-seq datasets. We calculated the communication probability for each ligand receptor pair across all samples and harmonized the dataset by imputing 0 communication probability for instances where a pair of ligand-receptor communication was not detected in a particular sample (but the exact ligand-receptor pair had > 0 % communication probability in another sample of the aggregate dataset). The communication probability for each ligand-receptor interaction was averaged for each distinct network/pathway observed in this study.

## Results

### Annotation of cell populations in the DTH biopsies in individuals with prior *Leishmania* infection

Paraffin embedded blocks of archived DTH biopsies obtained from individuals who reacted positive on LST were freshly sectioned and prepared for spatial transcriptomic analysis (supplementary Fig1A). The skin biopsy sections were then analyzed using sequencing and computational methods (Fig 1A). Following illumina sequencing, the quality of the sequencing data from the healthy sample and the five DTH biopsy samples was assessed by total read count (the number of RNA molecules detected per cell, Fig 1B), feature count (number of genes detected per cell, Fig 1C), the mitochondrial content (percentage of mitochondrial genes, which is a marker for cell stress or death, Fig 1D) and the sequence reads from hemoglobin (percentage of hemoglobin genes to measure for blood contamination, Fig 1E). Data showed a low percentage of mitochondrial content, near zero levels of hemoglobin across all samples indicating good cell viability. Overall, these plots demonstrate that high-quality data was obtained from all biopsies. UMAP of the pooled reads from all the five biopsies showed comparable distribution of cell populations (Fig 1F). Cell Ranger analysis followed by annotation tools in Seurat R package allowed clustering (Fig 1G) and annotation of cell types using marker genes characteristic of each cell type. Dot plots (Fig 1H-S) resolved the cell populations into 12 distinct clusters with minimal overlap.

**Figure 1.**
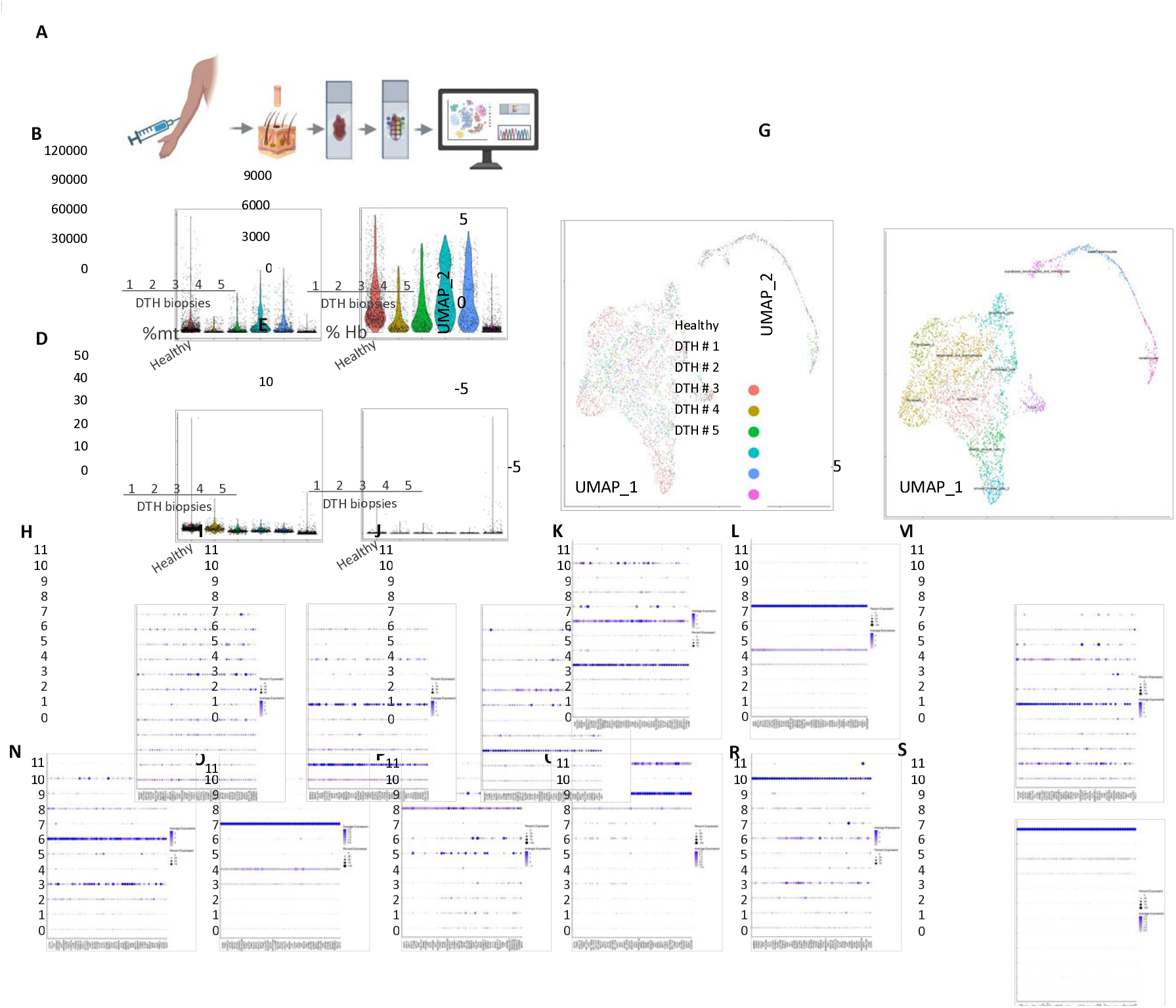
Spatial transcriptomic analysis reveals the distribution of cell types in control skin biopsy (Healthy) and biopsies of delayed type-hypersensitivity response site. A) Experimental design where residual FFPE tissues from DTH skin biopsies (n=5) and a control skin biopsy from a living donor (n=1) were used to perform scRNA-Seq using 10x Genomics Visium. B) Counts of the reads from illumina sequencing, C) feature counts, D) % of mitochondrial and E) % of hemoglobin transcripts are shown. F) UMAP plot from scRNA-Seq using Seurat package in R studio and the G) distribution of cells from Healthy and DTH biopsies into 12 clusters by unsupervised clustering is shown. H-S) Dot plots depicting the expression of top 50 marker genes used to characterize the clusters are shown.

### Spatial transcriptomic analysis reveals infiltration of immune cells in the DTH biopsies

After removing the damaged or dead cells and potential cell doublets, based on the %mt and %Hb read content, cell transcriptomes were retained for further analysis from the five skin biopsies as scRNAseq data. Mapping of the sequence reads to the H&E-stained sections of the skin biopsies revealed uniform distribution of the cells (spatial dim plots) from all the five biopsies (Fig 2A). A spatial resolution suitable for the identification of major cell types in the skin tissue including various structural cells such as fibroblasts, endothelial cells, keratinocytes, and immune cells such as Langerhans cells, macrophages was chosen (0.5). Accordingly, the UMAP of the identified cell types represented all the major cell types in human skin (Fig 2B). Transcriptomic identification of the cell types allowed a comparison of the cell composition in DTH biopsies compared to normal healthy skin (Fig 2C). A total of 1522 cells from the healthy biopsy and between 392 and 834 cells from each of the DTH biopsies were identified. Consistent with previous reports, an accumulation of immune cells in the DTH response site was observed (represented as percentages, Fig 2C). Compared to normal healthy skin that showed 4% of immune cells, DTH response site showed 6-38% of the cells identified as immune cells (Fig 2C). Enrichment of Langerhans cells was also noticeable in DTH tissue (up to 10.4%) compared to normal healthy tissue (3.2%, Fig 2C). Lymphatic endothelial cells and granulocytes in contrast showed a reduction in the DTH tissue (0.8-4.6%) compared to normal healthy tissue (6.1%) (Fig. 2C). Since the number of cells in the immune cell cluster were fewer (63-319), rather than subclustering the immune cells, we performed a de novo clustering of total cells in each biopsy to identify immune cells comprising of major cell types as macrophages, T and B cells, and granulocytes (Fig 2D). Cells in macrophages/basophil cluster were enriched in DTH tissue (25-63%) compared to normal healthy tissue that contained 9% of the cells belonging to this cluster (Fig 2D).

**Figure 2.**
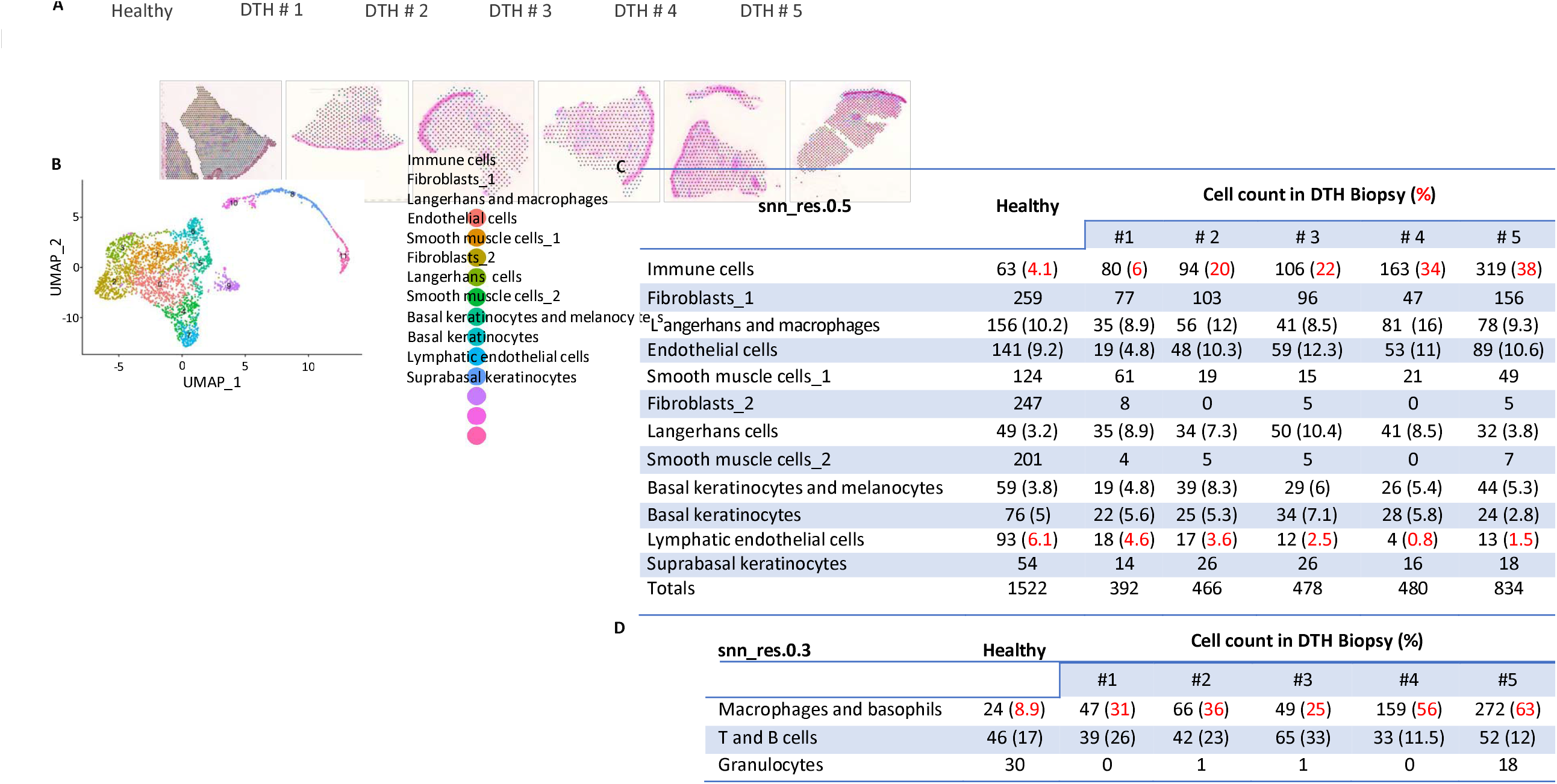
Spatial dim plots showing the organization of various cell types in healthy and DTH biopsies. A) Dim plots overlaid on the H&E stained sections of the healthy and DTH skin biopsies showing the spots sampled in each biopsy. B) Discrete cellular compositions identified based on marker gene expression are indicated in Healthy and DTH skin biopsies. C) The number of cells in each cluster detected in healthy and DTH skin biopsies is catalogued. C) The immune cell cluster is further decomposed to identify macrophages, T/B cells and granulocytes in the biopsies.

### DESeq analysis shows elevated transcriptional activities in immune cells, Langerhans and macrophages

As a first step towards gaining a mechanistic understanding of the DTH response, we assessed the transcriptional activities of cells in each of the 12 identified clusters. (Fig 3A-J). Significantly elevated transcriptional activities were observed in immune cells (Fig 3A), endothelial cells (Fig 3B), Langerhans and macrophages (Fig 3C), smooth muscle cells (Fig 3D), fibroblasts (Fig 3G), and lymphatic endothelial cells (Fig 3J) in DTH response compared to normal healthy skin. Due to the important role of immune cells in orchestrating the DTH response, we focused on the transcriptional activities in the clusters representing immune cells (Fig 3K) and Langerhans and macrophages (Fig 3L). A heatmap was generated showing the top 50 transcripts sorted on the basis of Log_2_Fc ratio between normal healthy tissue and the 5 DTH biopsies in immune cells identified transcripts such as type-I collagen (COL1A1, COL1A2), CXCL10, Cathepsin-K (CTSK), DCD and extracellular matrix protein such as tenascin-Xb (TNXB), all with reported roles in orchestrating DTH responses (Fig 3K). Similar analysis with cells in Langerhans and macrophage cluster showed enriched transcripts CD27, CD2, CD3D, LGALS1, CD247 and CXCL10 with well described roles in mediating DTH responses (Fig 3L).

**Figure 3.**
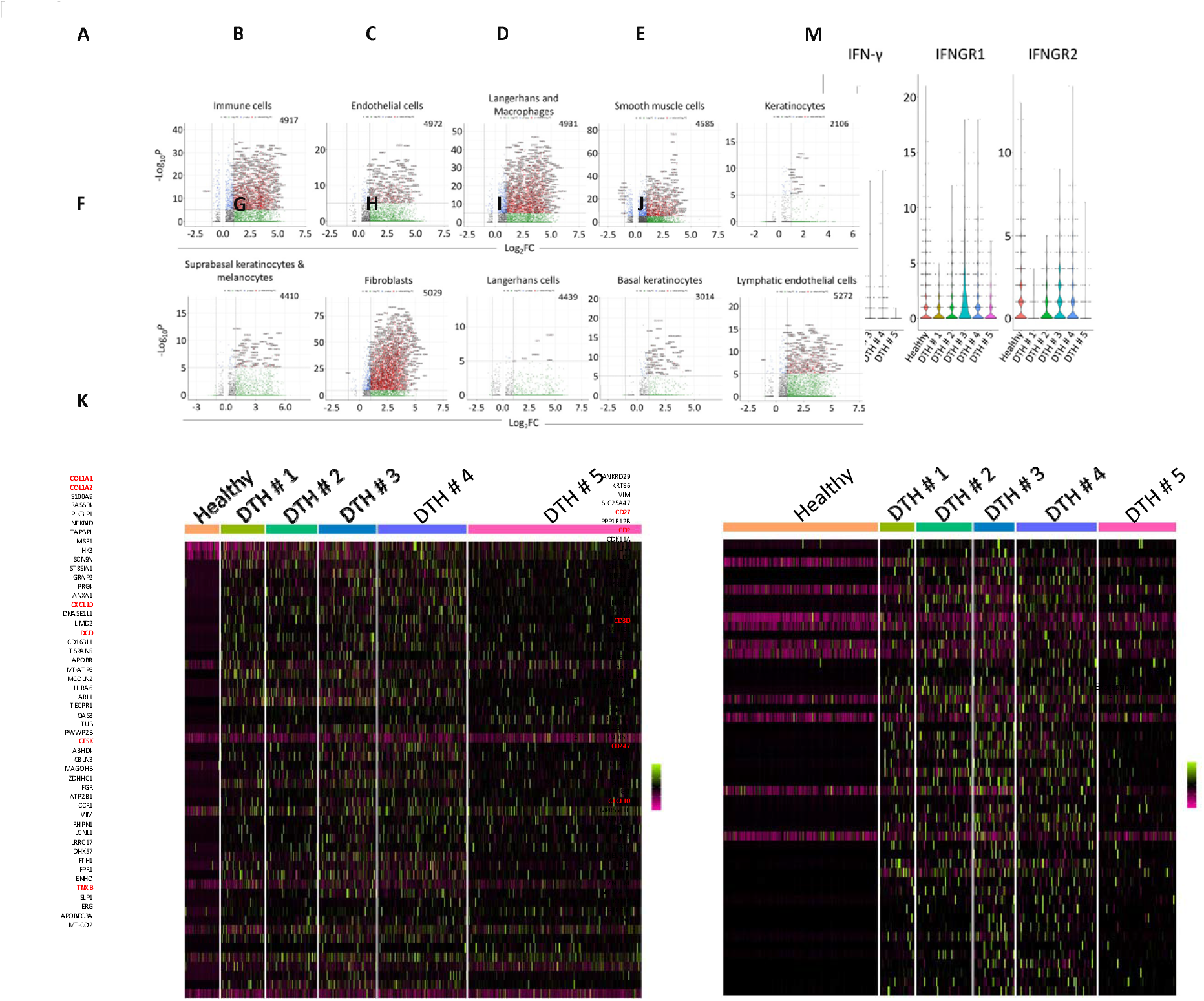
DESeq analysis among distinct cell clusters identified in healthy and DTH skin biopsies using spatial transcriptomic analysis. Transcriptionally active cell types in DTH skin biopsies compared to healthy skin biopsy are identified using DESeq analysis in A) immune cells, B) endothelial cells, C) Langerhans cells and macrophages, D) smooth muscle cells, E) keratinocytes, F) Suprabasal keratinocytes and melanocytes, G) fibroblasts, H) Langerhans cells, I) basal keratinocytes, and J) lymphatic endothelial cells. K) Heatmap of the top 50 genes sorted based on Log_2_Fc ratio in healthy and DTH biopsies in cells from the immune cell cluster and L) Langerhans cells and macrophage cluster are shown. M) The raw reads corresponding to IFN-γ and IFNGR1 and IFNGR2 transcripts detected in healthy and DTH skin biopsies are shown.

### Cell communication network analysis shows IL-16 and TNF signaling in Langerhans cells

Towards developing a mechanistic understanding of various cell types in the DTH response, we constructed cell to cell communication network among 12 distinct clusters. Due to the known roles of IL-16 and TNF in recruiting CD4^+^ T cells and leukocytes respectively in previous studies of DTH response, we first focused on IL-16 and TNF. Violin plots of the IL-16 expression among 12 clusters showed that clusters 1 (Fibroblasts), 2 (Langerhans cells and macrophages), 6 (Langerhans cells), 9 (Basal Keratinocytes), and 10 (Lymphatic endothelial cells) were the major producers of IL-16 in DTH biopsies compared to healthy tissue that showed baseline levels of IL-16 (Fig 4A-B). To visualize the expression on IL-16, dim plots of the five DTH and normal healthy biopsies were constructed that showed spatial location of the cell types that produced IL-16 (Fig 4C). Of the various cell types producing IL-16, the communication probabilities between cell clusters showed that Langerhans cells as the main senders and other immune cells such as endothelial cells, immune cells, LECs and other structural cells such as smooth muscle cells, Suprabasal keratinocytes and melanocytes and fibroblasts as the recipients were enriched in the DTH tissue compared to healthy tissue (Fig 4D). Similarly, violin plots of TNF expression among 12 clusters showed that clusters 5 (Fibroblasts_2), 6 (Langerhans cells), 8 (Basal keratinocytes and melanocytes), and 11 (Suprabasal keratinocytes) were the major producers of TNF in DTH biopsies compared to healthy tissue that showed baseline levels of TNF and its receptor TNFRSF1B (Fig 4E-H). To visualize the expression on TNF, dim plots of the five DTH and normal healthy biopsies were constructed that showed spatial location of the cell types that produced TNF (Fig 4I). Of the various cell types producing TNF, the communication probabilities between cell clusters showed that Langerhans cells as the main senders and other immune cells such as endothelial cells, immune cells, LECs and multiple other structural cells such as smooth muscle cells, basal keratinocytes, suprabasal keratinocytes and fibroblasts as the recipients were enriched in the DTH tissue compared to healthy tissue (Fig 4J).

**Figure 4.**
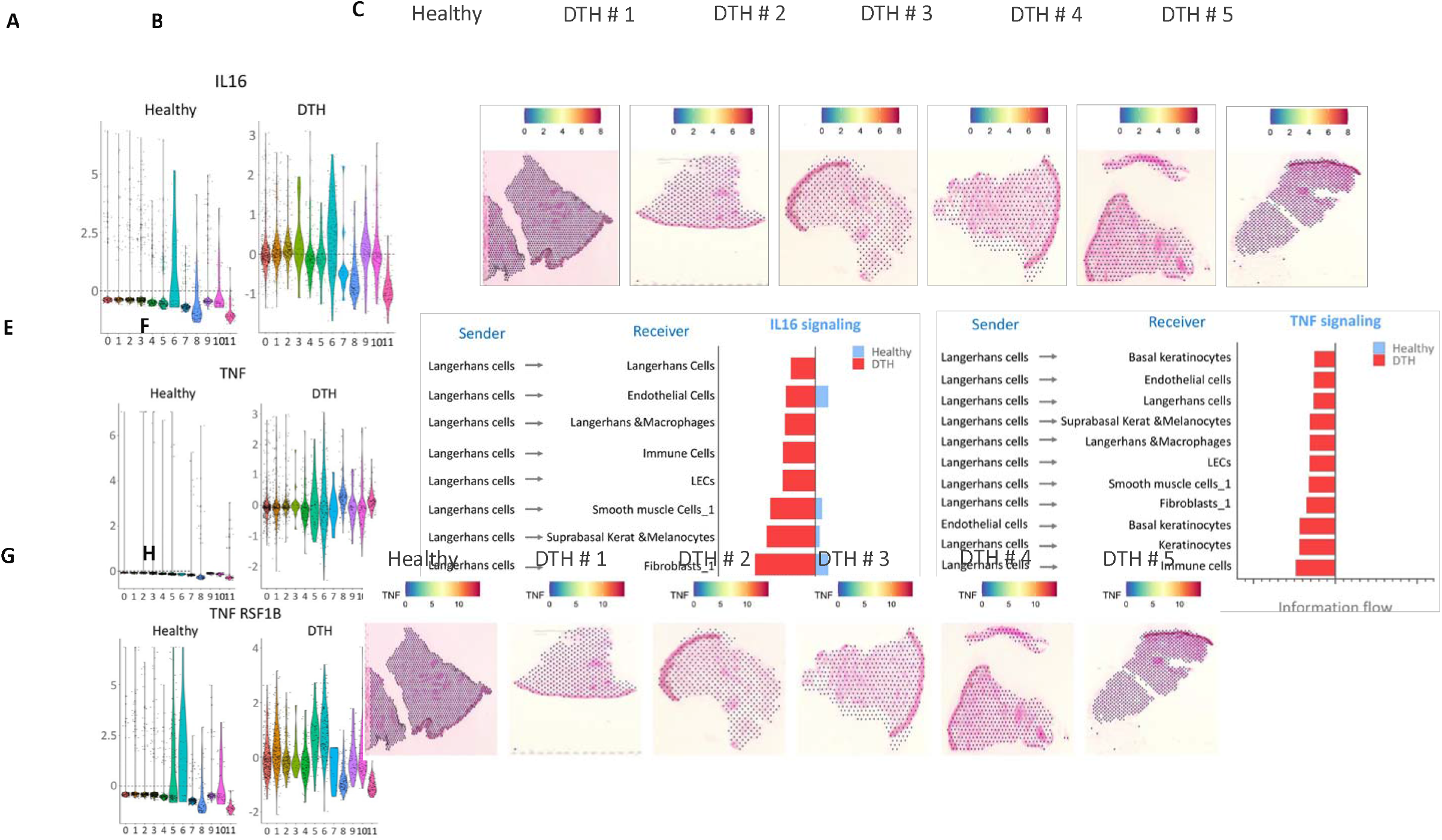
IL-16 and TNF signaling is enriched in DTH biopsies. A) Violin plots showing the reads of IL-16 transcripts in various cell types from healthy and B) DTH biopsies. C) Dim plots overlaid on the H&E stained sections of the healthy and DTH skin biopsies showing the IL16 transcripts in each biopsy. D) Communication networks based on the observed absolute differences in the calculated communication probabilities between healthy and DTH biopsies are graphed to highlight the communication between Langerhans cells and various receivers. E) Violin plots showing the reads of TNF transcripts in various cell types from healthy and F) DTH biopsies. G) Violin plots showing the reads of TNF receptor TNFRSF1B transcripts in various cell types from healthy and H) DTH biopsies. I) Dim plots overlaid on the H&E stained sections of the healthy and DTH skin biopsies showing the TNF transcripts in each biopsy. J) Communication networks based on the observed absolute differences in the calculated communication probabilities between healthy and DTH biopsies are graphed to highlight the communication between Langerhans cells and various receivers.

### Chemokines CCL2, CCL8 and CCL19 are highly enriched in DTH tissue

To delineate the role of chemokines in cell recruitment events intricately associated with the DTH response, we analyzed the expression of major chemokines in all the 12 clusters. Normalized transformed read counts of chemokines CCL2, CCL5, CCL8, CCL13, CCL19, CCL21 and CCL22 were plotted across all 12 clusters (Fig 5A-G). Results showed that CCL2, CCL8 and CCL19 were noticeably elevated in the DTH tissues (Fig. 5 A, C, and E) compared to normal healthy tissue in most of the clusters and CCL5, CCL13, CCL21 and CCL22 similarly showed an elevated expression though to a lesser degree in the DTH tissues compared to normal healthy tissue (Fig 5 B,D,F, G). Cell clusters are color coded and indicated in Fig 5H. Dot plots of the average of normalized and transformed counts of the selected chemokine transcripts detected in Langerhans and macrophage cluster are shown highlighting the role of these cells in secreting and orchestrating the DTH response (Fig 5I). Dim plots of CCL2, CCL8 and CCL19 showing the expression of these ligands across all cell types are included to illustrate tαhe spatial distribution of the transcripts of these CCLs (Fig 5J-L).

**Figure 5.**
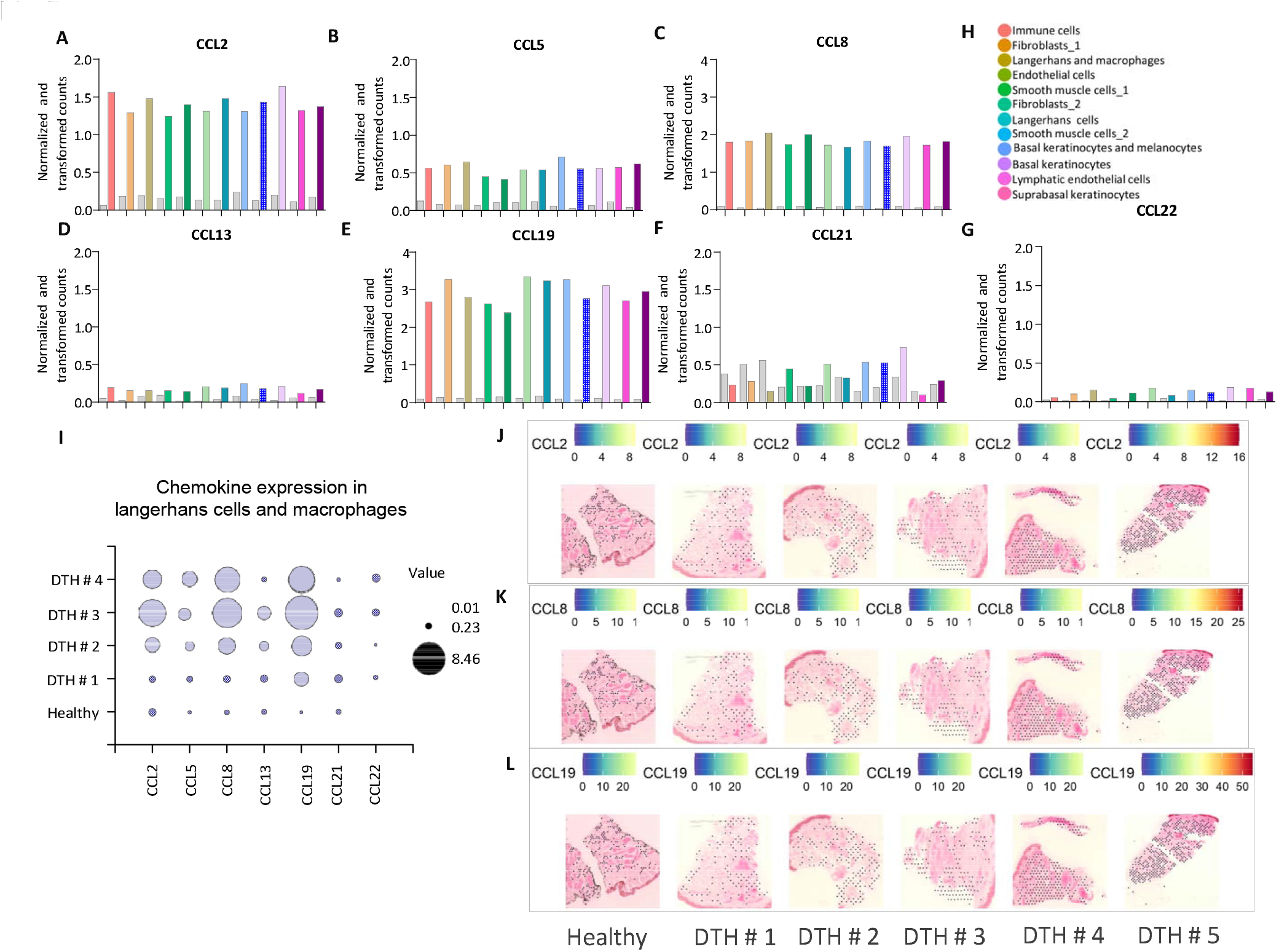
Chemokines CCL2, CCL8, and CCL19 are highly enriched in DTH biopsies. Normalized and transformed counts of chemokines A) CCL2, B) CCL5, C) CCL8, D) CCL13, E) CCL19, F) CCL21 and G) CCL22 in each cell type identified are shown. H) Map of the cell types identified based on marker gene expression in Healthy and DTH skin biopsies is color coded. I) Bubble chart showing the expression of various cytokines in Langerhans and macrophage cell cluster as an indication of the significant contribution from this cell type. Dim plots overlaid on the H&E stained sections of the healthy and DTH skin biopsies showing the J) CCL2, K) CCL8, and L) CCL19 transcripts in each biopsy.

### CCL19-CCR7 signaling is the most significant signaling observed in DTH response

Further analysis of the expression of CCLs and their receptors revealed that CCL2, CCL8 and CCL19 are widely expressed in all the clusters (Fig 6A). More importantly, the chemokine receptor CCR7 was expressed mainly in Fibroblasts_1, Fibroblasts_2, Langerhans cells, and Basal Keratinocytes (Fig 6A). Chemokine receptors CCR1 was found mainly expressed in Fibrobalsts_1, Langerhans cells and macrophages, Smooth muscle cells_1, Fibroblasta_2, Langerhans cells and Basal Keratinocytes where as CCR2 was mainly expressed in Fibroblasts_2, Langerhans cells and Smooth muscle cells_2 (Fig 6B-C). The strength of communication as shown in the heatmap indicated that Langerhans cells are the major contributors to the chemokine signaling among all other cell pairs, followed by Fibroblasts, Basal keratinocytes and LECs (Fig 6C). Chord diagrams of the chemokine expression showed that Langerhans cells and endothelial cells (indicated by the arc length) play important roles in this signaling (Fig 6D). When the relative contribution of the observed ligand-receptor pairs was sorted, CCL19-CCR7 interaction stood out as the most enriched followed by CCL8-CCR1, CCL8-CCR2 and others in decreasing order (Fig 6E). Chord diagrams constructed to delineate the cell types involved in chemokine signaling revealed that CCL19-CCR7 interaction mainly occurred between Langerhans cells (purple arcs), endothelial cells and LECs (Fig 6F). Similarly, CCL8-CCR1 interaction occurred mainly between Langerhans cells, endothelial cells, Keratinocytes, Langerhans cells macrophages, immune cells, LECs and smooth muscle cells (Fig 6G). CCL2-CCR2 interaction occurred mainly between smooth muscle cells_2, Langerhans cells, endothelial cells, Keratinocytes, Langerhans cells macrophages, immune cells, and LECs (Fig 6H). CCL5-CCR1 interaction on the other hand occurred mainly between Langerhans cells, endothelial cells, smooth muscle cells_1, Langerhans cells, endothelial cells, Keratinocytes, Langerhans cells macrophages, immune cells, and LECs (Fig 6I).

**Figure 6.**
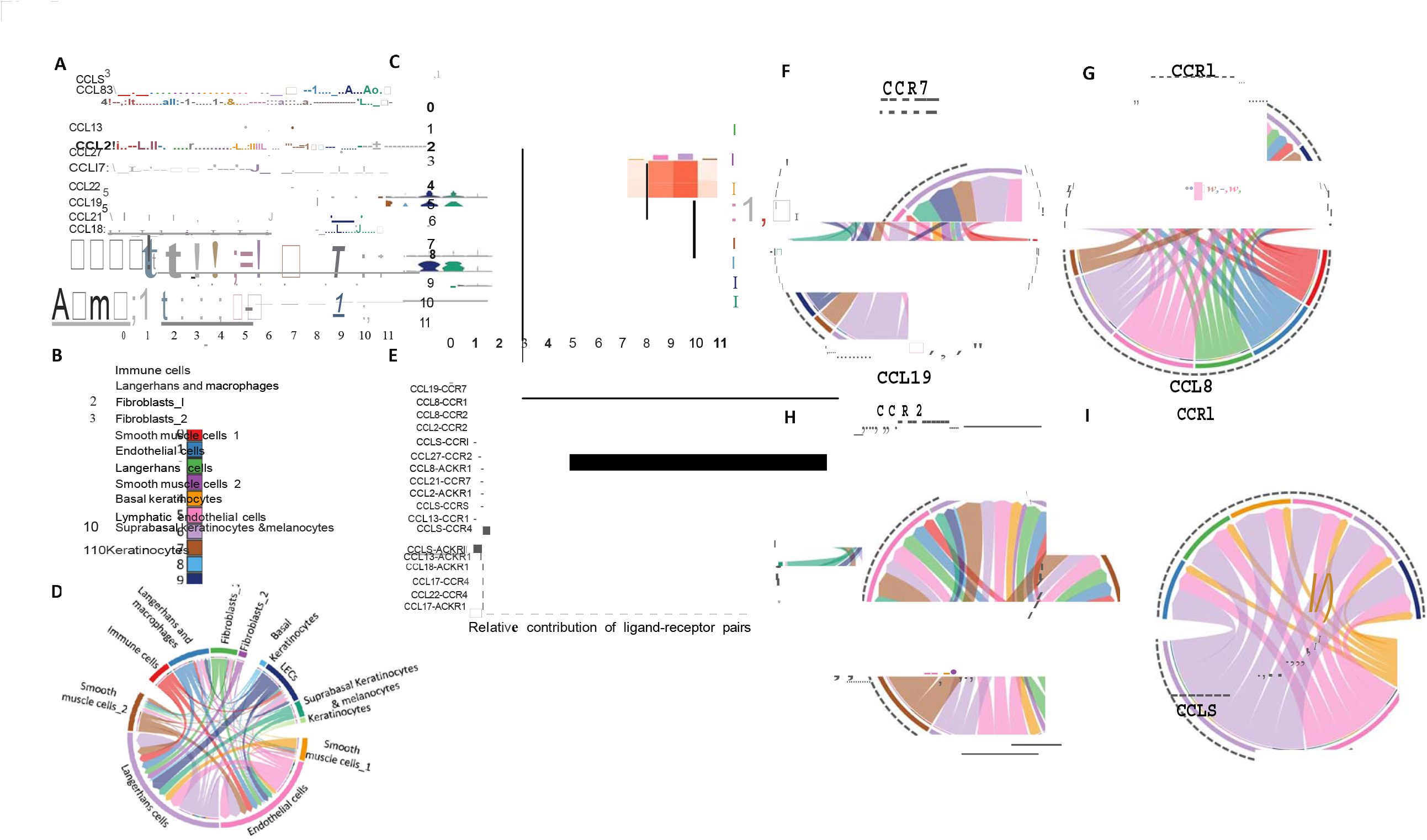
Chemokine communication network among cell types in DTH reaction. A) Violin plots of various chemokines detected in each cell type are shown. B) Color coded map of the cell types identified based on marker gene expression in healthy and DTH skin biopsies. C) Heatmap of the chemokine signaling observed among senders (x-axis) and receivers (y-axis) with the numbers representing cell types, Langerhans cells being the predominant mediators. D) Chord diagrams showing the predominant role of Langerhans cells and endothelial cells in addition to other cell types in chemokine signaling in DTH reactions. E) Bar graph showing the relative contribution of ligand-receptor pairs with CCL19-CCR7 as the most dominant signaling. Chord diagrams showing the communication between various cell types with various ligand-receptor pairs, i.e., F) CCL19-CCR7, G) CCL8-CCR1, H) CCL2-CCR2, and CCL5-CCR1 in DTH biopsies.

## Discussion

In our spatial transcriptomics and functional studies, we identified multiple cytokine and chemokine systems that work in concert to orchestrate delayed-type hypersensitivity (DTH) reactions in skin, each contributing distinct but complementary functions. IL-16 serves as the primary temporal orchestrator of DTH responses. This CD4^+^ T cell chemoattractant was detected within 6 hours post-antigen challenge and increases rapidly with tissue swelling. IL-16 functions through both direct recruitment of antigen-specific Th1 cells and indirect amplification of inflammatory cascades, including upregulation of secondary mediators like MIP-1α. Neutralization studies demonstrate that IL-16 is functionally significant, reducing DTH responses by approximately 50% and decreasing infiltration of multiple leukocyte populations, demonstrating the central role of IL16 In DTH response (Yoshimoto 2000). Elevated levels of IL-16 in the DTH biopsies in our spatial transcriptomic analysis is thus consistent with the data supporting its role in DTH reactions against model antigens such as methylated BSA (Yoshimoto 2000).

The role of IL-8 in recruiting neutrophils to the DTH response has been demonstrated in tuberculin skin reactions (Larsen et al., 1995). The tuberculin (Mantoux) reaction response is accompanied by a prominent local chemokine/cytokine milieu and an early granulocyte component; blockade of IL-8 markedly suppresses the tuberculin skin reaction and reduces neutrophil influx (Larsen et al., 1995). Similarly, immunohistochemical analysis of human DTH biopsies showed the presence of cytokines IL-lα, IL-lβ, IL-6, IFN-γ and TNF-α (Chu et al., 1992, Tsicopoulos, et al., 1992). Absence of IL-17 diminished the DTH response to PPD and also reduced IFN-γ production (Umemura 2007). In leishmanin skin tests, IL-17 was reported mainly in PBMCs of LST positive individuals (Bahrami et al., 2014). Since our analysis involved sampling of fully developed DTH response site, neutrophil mediated IL-17 production was not detected and granulocytes as a whole were not detected in significant numbers in the DTH biopsies (Fig 2D, supplementary fig 2). Chemokines CXCL1, 2, 5 and 8 that act downstream of IL-17 similarly were not detected in our DTH biopsies (supplementary fig 2) further suggesting that the synergistic effects of IL-17 and TNF in neutrophil recruitment may not be evident in the effector phase of the DTH response in leishmanin skin tests.

To delineate the drivers of DTH response in leishmanin skin tests, we focused on chemokine signaling networks. Our spatial transcriptomic studies revealed that CCL19-expressing COL18A1+ fibroblasts interact closely with CCR7+ LAMP3+ dendritic cells in leukocyte-infiltrated areas, creating organized microenvironments resembling tertiary lymphoid structures. This axis appears critical for optimal antigen presentation through selective recruitment of mature dendritic cells and spatial coordination of DC-T cell interactions. CCL19-CCR7 signaling functions as the spatial organizer of immune responses and has been mainly studied in lymph node organization (Forster et al., 1999, 2008). Similar roles for orchestrating local immune response in skin for CCL19-CCR7 has been reported in systemic sclerosis and psoriasis studies (Mathes et al., 2014, Lee et al., 2024).

The chemokine receptor CCR7, with its principal ligand CCL19 (and CCL21), is critical in orchestrating multiple phases of the immune response. These chemokines are homeostatically expressed in secondary lymphoid organs, particularly within T cell zones, high endothelial venules, and lymphatic endothelium. CCR7 is expressed on naïve and central memory T cells, as well as on mature dendritic cells (DCs) (Muller 2002). In the context of delayed-type hypersensitivity, CCR7–CCL19 may facilitate the initiation phase by ensuring that antigen-loaded DCs migrate from peripheral tissue into draining lymph nodes, and that naïve antigen-specific T cells home to T cell zones for priming. Disruption of CCR7 or its ligands results in disorganized lymphoid architecture, impaired DC and T cell localization, and severely attenuated DTH responses, as illustrated by CCR7-deficient mice lacking proper contact sensitivity and DTH reactions (Muller 2002).

Effector memory T (Tem) cells which mediate the elicitation phase of DTH are thought to rely less on CCR7 for tissue entry (since CCR7 is downregulated), but CCR7-mediated steps upstream (priming, central memory maintenance) strongly influence the magnitude and quality of the effector phase. Imaging studies in DTH models show that CCR7-negative effector T cells infiltrate challenge sites and interact with antigen-presenting cells, underscoring that although CCR7 is less used at late stages, its prior signaling is essential for generating these effectors (Matheu 2008, Kuwabara 2009).

Additionally, CCR7–CCL19 signaling also appears to affect the resolution phase: CCR7 expression allows activated T cells and DCs to exit tissues via lymphatics, contributing to the waning of inflammation. Loss of CCR7 signaling is associated with prolonged or altered immune activation (Debes et al., 2005). While direct studies on CCL19’s unique contribution (versus CCL21) in DTH remain relatively limited, analogous evidence from contact hypersensitivity and other Th1/Th17 models support nonredundant roles of its ligands in shaping both the priming and polarization of T cell responses.

In addition to IL-16 in T cell recruitment to the DTH site (Cruikshank et al., 2000), CCL19-CCR7 may act as principal mediators of DTH response. Our DESeq analysis revealed potential roles of several other immunomodulators such as Cathpesin-K, Collagen, AnnexinA1 and TenascinX. Cathepsin K is a protease expressed notably in osteoclasts but also in macrophages and some antigen-presenting cells; it degrades extracellular matrix components (e.g. collagen) and participates in antigen processing. In DTH, ECM degradation is important for immune cell infiltration and for remodeling of tissue during lesion formation. Cathepsin K activity could facilitate migration of macrophages and lymphocytes by breaking down collagen and also influence antigen presentation via peptide generation (Paracha 2022). While specific studies on Cathepsin K in DTH are limited, other cathepsins (e.g. CatS), there is evidence from inflammatory models that CatK contributes to tissue damage and remodeling; inhibition of its activity attenuates certain Th1-mediated pathologies (Schwenk 2019).

TNX is an extracellular matrix glycoprotein involved in ECM structure, particularly collagen fibrillogenesis, binding to collagens and tropoelastin, and stabilizing ECM architecture (Egging 2007). In DTH responses, where ECM breakdown and subsequent repair/remodeling occur, TNX may influence how well collagen fibrils are re-assembled, how stable the ECM scaffold is, and thus how induration, fibrosis or resolution proceeds. For instance, TNX deficiency results in reduced collagen deposition, skin fragility, altered elasticity. Accordingly, Tnxb^-/-^ mice show reduced dermal collagen content/strength and broad ECM abnormalities, mechanistically relevant to DTH induration/remodeling but no direct CHS/DTH challenge study (Mao 2002).

While the TNF and IL-16 detected in our data likely contribute as an amplifying mediator that may be non-essential due to pathway redundancy. While TNF family receptors participate in immune cell interactions and inflammatory cascades, the presence of multiple mechanisms suggests DTH can proceed through alternative pathways. Whereas IL-16 provides broad temporal coordination and initial cell recruitment, CCL19-CCR7 ensures spatial organization and functional efficiency. Integrated analysis undertaken herein revealed the multi-layered system that mediates inflammatory immune responses in the skin. Collectively, our spatial transcriptomic analysis revealed the central role of IL-16 and CCL19-CCR7 signaling in the DTH response induced by inoculation of leishmanin antigens. More importantly our unbiased analysis revealed unexpected roles for other immunomodulators such as cathepsin K, Collagen A, and TenascinX proteins. The mechanistic roles of these proteins in DTH response will be formally demonstrated in future studies. Our data supports the development of commercial scale leishmanin antigens by identifying product standards that may be examined in clinical trials of leishmanin test antigens in endemic areas.

## Acknowledgements

We thank Spyros Karaiskos and Nikki Tirrell for their bioinformatics support and help with data analysis

## Figure legends

**Supplementary figure 1:**
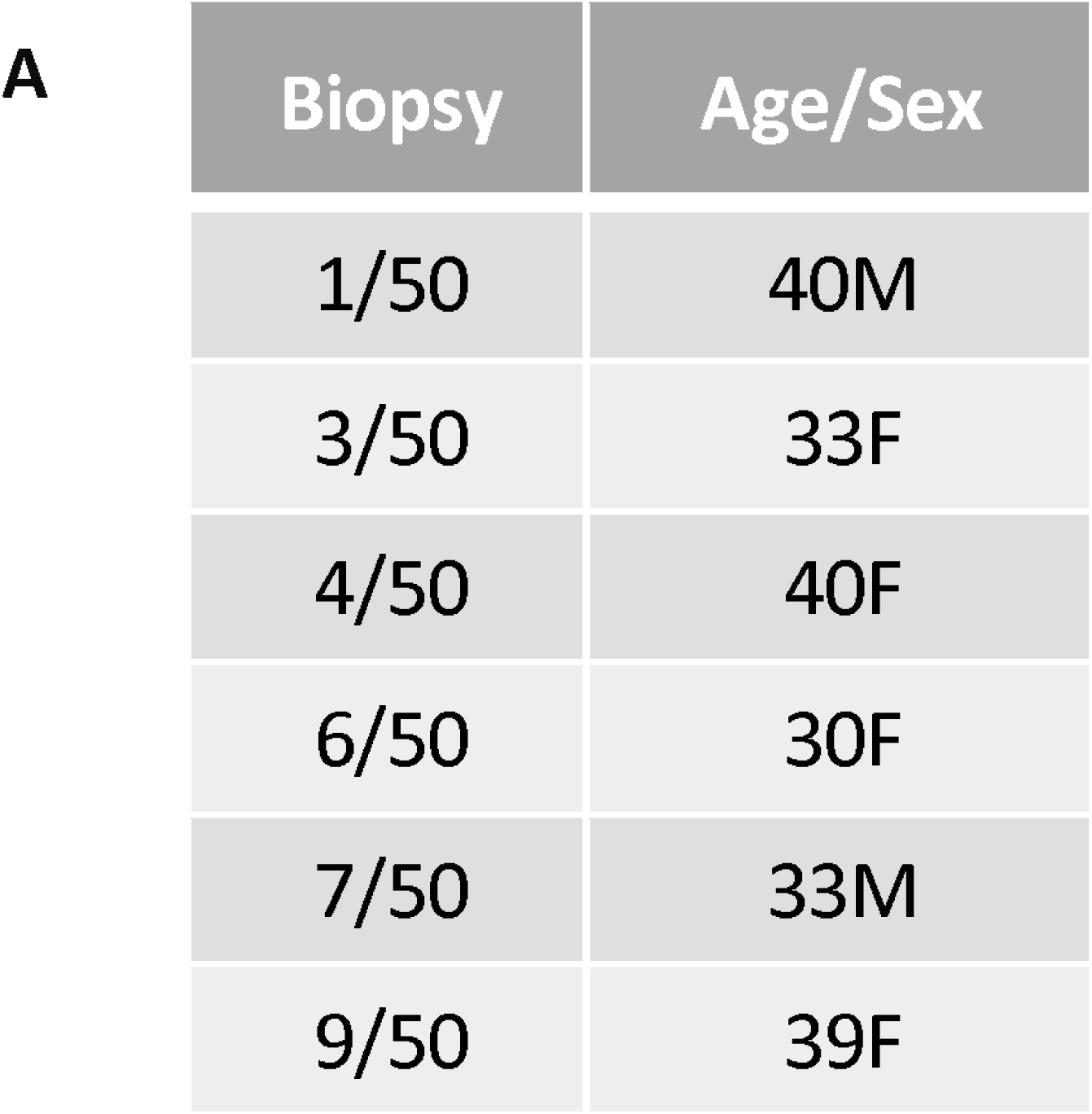
Age/Sex of the LST positive and negative subjects

**Supplementary figure 2:**
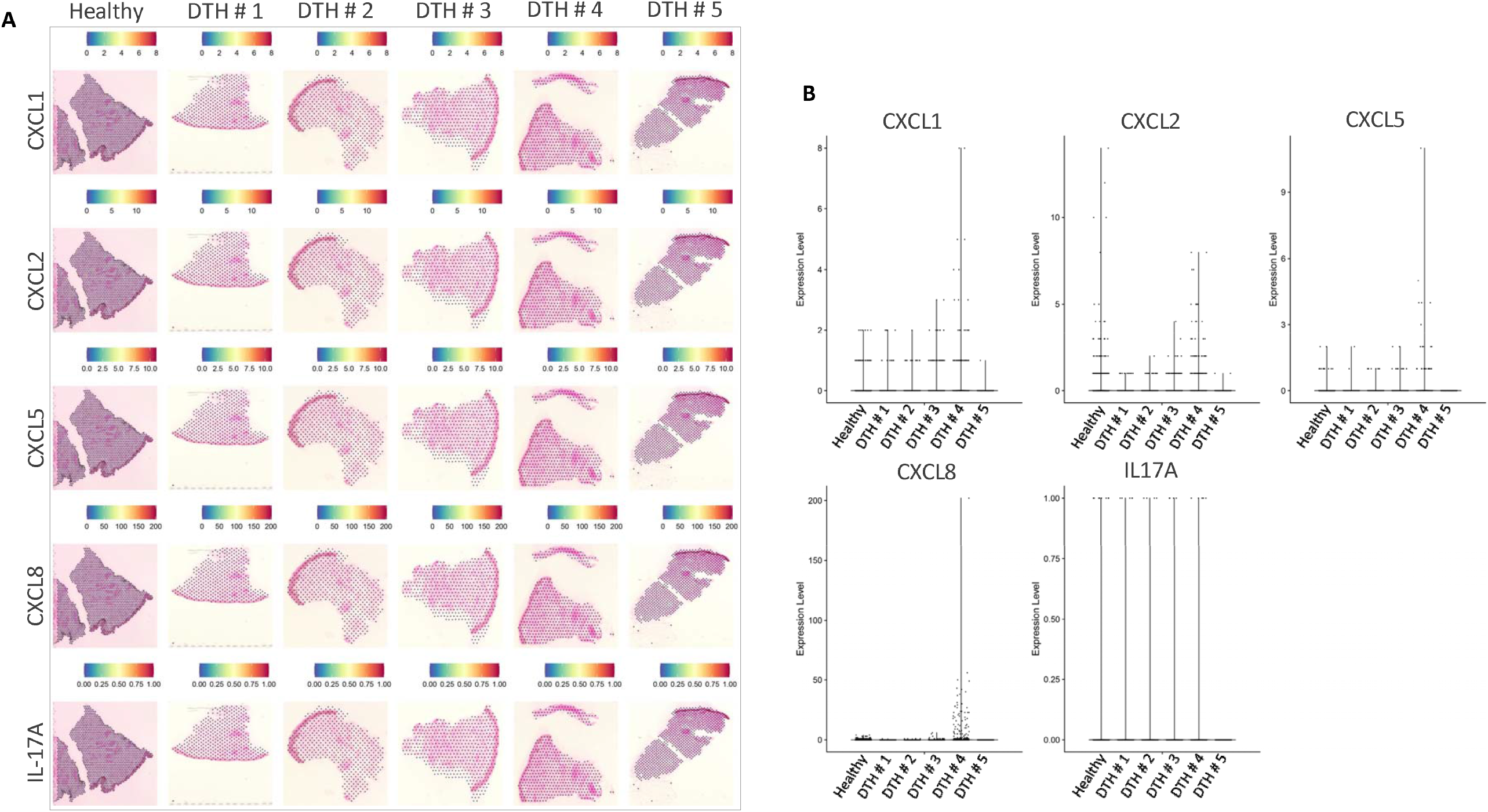
IL-17 and the downstream chemokines are not detected in DTH biopsies. A) Dim plots of the healthy and DTH biopsies showing the expression of CXCL1, CXCL2, CXCL5, CXCL8 and IL-17A. B) Violin plots for the same chemokines in each biopsy are also shown.

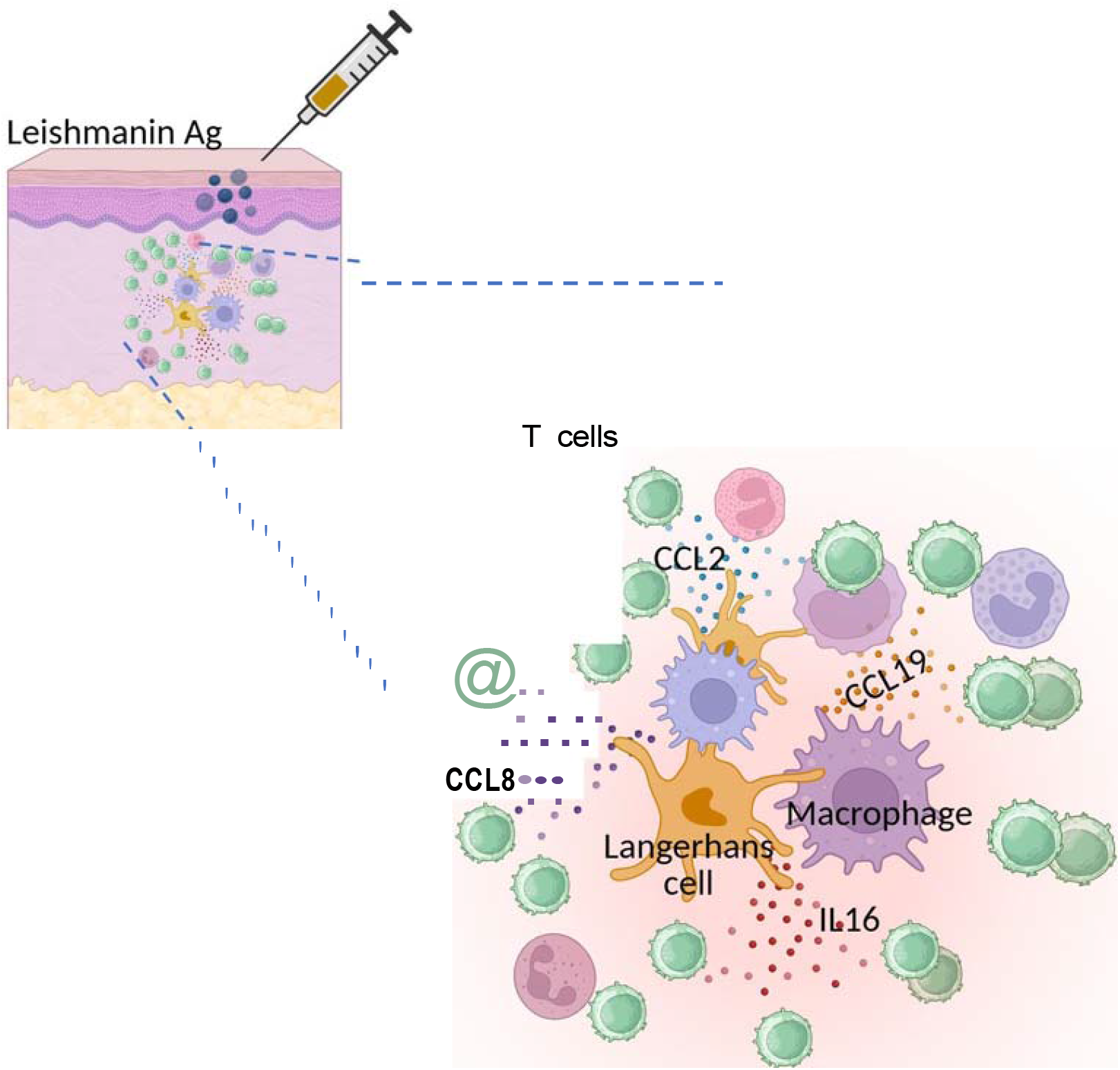

## Notes

### Competing Interest Statement

The authors have declared no competing interest.

### Funding

- Global Health Innovative Technology Fund

